# The hallucinogenic serotonin_2A_ receptor agonist, DOI, promotes CREB-dependent gene expression of specific plasticity-associated genes in the rodent neocortex

**DOI:** 10.1101/2021.10.19.464691

**Authors:** Lynette A. Desouza, Madhurima Benekareddy, Sashaina E. Fanibunda, Farhan Mohammad, Tamar Gur, Julie A. Blendy, Vidita A. Vaidya

**Affiliations:** Department of Biological Sciences, Tata Institute of Fundamental Research, Mumbai 400005, India; Medical Research Centre, Kasturba Health Society, Mumbai 400056, India; Department of Psychiatry and Behavioral Health, The Ohio State University College of Medicine, Columbus, OH, 43210, United States; Department of Systems Pharmacology and Translational Therapeutics, Perelman School of Medicine, University of Pennsylvania, Philadelphia, PA 19104, United States

**Keywords:** 5-HT_2A_ receptor, cAMP response element binding protein, immediate early gene, BDNF, Arc, serotonergic psychedelic, cortical neuron, CREB deficient mice

## Abstract

Psychedelic compounds that target the 5-HT_2A_ receptor are reported to evoke psychoplastogenic effects, including enhanced dendritic arborization and synaptogenesis. Transcriptional regulation of neuronal plasticity-associated genes is implicated in the cytoarchitectural effects of serotonergic psychedelics, however the transcription factors that drive this regulation are poorly elucidated. Here, we addressed the contribution of the transcription factor cAMP response element binding protein (CREB) in the regulation of neuronal plasticity-associated genes by the hallucinogenic 5-HT_2A_ receptor agonist, DOI. *In vitro* studies with rat cortical neurons indicated that DOI enhances the phosphorylation of CREB (pCREB) through the MAP kinase and CaMKII pathways, with both cascades contributing to the DOI-evoked upregulation of *Arc, Bdnf1, Cebpb* and *Egr2* expression, whilst the upregulation of *Egr1* and *cFos* mRNA involved the MAP kinase and CaMKII pathway respectively. We observed a robust DOI-evoked increase in the expression of several neuronal plasticity-associated genes in the rat neocortex *in vivo*. Further, 5-HT_2A_ receptor stimulation enhanced pCREB enrichment at putative cAMP response element (CRE) binding sites in the *Arc, Bdnf1, Cebpb, cFos*, but not *Egr1* and *Egr2*, promoters in the rodent neocortex. The DOI-mediated transcriptional induction of *Arc, cFos* and *Cebpb* was significantly attenuated in the neocortex of CREB deficient (CREBαδ KO) mice. Collectively, these results indicate that the hallucinogenic 5-HT_2A_ receptor agonist DOI leads to a rapid transcriptional upregulation of several neuronal plasticity-associated genes, with a subset of them exhibiting a CREB-dependent regulation. Our findings raise the intriguing possibility that similar to slow-acting classical antidepressants, rapid-action serotonergic psychedelics that target the 5-HT_2A_ receptor may also recruit the transcription factor CREB to enhance the expression of neuronal plasticity-associated genes in the neocortex, which could in turn contribute to the rapid psychoplastogenic changes evoked by these compounds.

## 1. Introduction

There has been a renewal of interest in serotonergic psychedelics as potential rapid-acting antidepressants for the treatment of anxiety and depression (Nutt et al., 2020; Vollenweider and Preller, 2020; Banks et al., 2021; De Gregorio et al., 2021). Most serotonergic psychedelics target the serotonin_2A_ (5-HT_2A_) receptor, and agonist action at the 5-HT_2A_ receptor is implicated in the molecular, cytoarchitectural, hallucinogenic and mood-related behavioral effects of serotonergic psychedelics (Vollenweider et al., 1998; González-Maeso et al., 2003, 2007; Martin and Nichols, 2016; López-Giménez and González-Maeso, 2018; Preller et al., 2018; Madsen et al., 2019). Diverse serotonergic psychedelics, as well as the hallucinogenic 5-HT_2A_ receptor agonist, 2,5-dimethoxy-4-iodoamphetamine (DOI), can elicit both unique and distinctive molecular, behavioral and electrophysiological effects (Aghajanian and Marek, 1999; González-Maeso et al., 2003, 2007; Marek, 2017; Banerjee and Vaidya, 2020). Common to all of these 5-HT_2A_ receptor agonists is a regulation of neuronal structural plasticity in the neocortex; from increased dendritic complexity to enhanced synaptogenesis (Ly et al., 2018, 2020; Savalia et al., 2021; Shao et al., 2021; Vargas et al., 2021). The rapid transcriptional effects evoked via agonistic action at the 5-HT_2A_ receptor, in particular the enhanced expression of neuronal plasticity-associated gene expression, are implicated in contributing to these psychoplastogenic effects (Nichols et al., 2003; Olson, 2018; Berthoux et al., 2019; Artin et al., 2021; de Vos et al., 2021; Jefsen et al., 2021).

5-HT_2A_ receptors are G-protein coupled receptors that drive Gq-signaling to activate phospholipase C beta (PLCβ) -mediated cleavage of phosphatidylinositol bisphosphate (PIP2) into inositol triphosphate (IP3) and diacylglycerol (DAG) (Raymond et al., 2001; Sharp and Barnes, 2020). IP3 mobilizes release of calcium from intracellular endoplasmic reticulum stores, which activates calcium/calmodulin dependent kinase II (CaMKII), while DAG activates protein kinase C (PKC) that phosphorylates and activates mitogen-activated protein (MAP) kinase (MAPK) signaling (Banerjee and Vaidya, 2020). These kinases in turn are reported to enhance the phosphorylation of the transcription factor, cyclic AMP response-element binding protein (CREB), that has been previously shown to regulate the expression of several neuronal plasticity-associated genes (Lonze and Ginty, 2002; Sakamoto et al., 2011; Belgacem and Borodinsky, 2017). CREB is also a central target for diverse classes of antidepressant treatments, contributes to the neurotrophic, neurogenic and behavioral effects of antidepressants (Nibuya et al., 1996; Duman RS, Nibuya M, 1997; Thome et al., 2000; Chen et al., 2001; Nair and Vaidya, 2006), and is dysregulated in both animal models of depression (Carlezon et al., 2005; Duman RS, 2005; Krishnan and Nestler, 2008, 2010) and in major depressive disorder patients (Blendy et al., 1996; Koch et al., 2009). Here, we sought to address whether CREB plays an important role in the regulation of neuronal plasticity-associated gene expression within the neocortex that arise in response to treatment with the hallucinogenic 5-HT_2A_ receptor agonist, DOI. Using both *in vitro* and *in vivo* approaches, as well as a CREB-deficient mouse line, we demonstrate that CREB contributes to the DOI-mediated regulation of a subset of neuronal plasticity-associated genes within the neocortex. This raises the intriguing possibility that similar to classical antidepressants, serotonergic psychedelics that target the 5-HT_2A_ receptor may also recruit CREB-mediated transcriptional regulation of specific neuronal plasticity-associated genes in the neocortex, thus contributing to the effects on neuronal structural plasticity and mood-related behaviour.

## 2. Materials and Methods

### 2.1 Animal treatments

Male Sprague-Dawley rats (2-3 months) bred in the Tata Institute of Fundamental Research (TIFR) animal facility and CREB deficient (CREBαδ KO) mice bred in the University of Pennsylvania animal facility were used for all experiments. For experiments using CREB deficient mice, the hypomorphic CREBαδ knockout mouse line that lacks the α and δ isoforms of CREB (CREBαδ KO) were used (Blendy et al., 1996). CREBαδ KO mice were generated as previously described (Walters and Blendy, 2001) and were maintained as F1 hybrids of 129SvEvTac:C57BL/6. For all experiments, CREBαδ KO mice and the wild type controls were generated via crossing heterozygote CREBαδ 129SvEvTac with heterozygote CREBαδ C57BL/6 mice, allowing for a uniform genetic background in the experimental cohort. For the establishment of *in vitro* cortical cultures, rat embryos were derived at embryonic day 17.5 (E17.5) from timed pregnant Sprague-Dawley dams. All animal procedures using Sprague-Dawley rats were carried out in accordance with the Committee for Care and Supervision of Experimental Animals (CPCSEA) and approved by the TIFR Institutional Animal Ethics Committee. All experiments with the CREBαδ KO mouse line were carried out in accordance with the NIH guideline for the care and use of laboratory animals and were approved by the University of Pennsylvania Animal Care and Use Committee. Animals were group housed and maintained on a 12 h light–dark cycle (lights on at 7 am) with access to food and water *ad libitum*.

Sprague-Dawley rats, CREBαδ KO and litter-matched wild type (WT) mice, received intraperitoneal injections of the 5-HT_2A_ agonist, 2,5-dimethoxy-4-iodoamphetamine (DOI, 8 mg/kg, Sigma-Aldrich, USA) or vehicle (0.9% NaCl) and were sacrificed two hours post drug administration.

### 2.2 Cortical neuron cultures

Primary cortical neuron cultures were established from E17.5 rat embryos as described previously (Desouza et al., 2011; Fanibunda et al., 2019). Rat embryonic cortices were dissected and treated with trypsin-EDTA for ten minutes, prior to dissociation in culture medium - Neurobasal medium supplemented with 2% B27 supplement, 0.5 mM L-glutamine, 5 U/ml penicillin and 5 U/ml streptomycin (Invitrogen, USA). Cells were plated on poly-D-lysine (Sigma-Aldrich, USA) coated 35 mm dishes at a density of 10^6^ cells/dish. Following attachment and neurite extension *in vitro* for a period of seven days, neurons were treated with DOI (10 μM) or vehicle (DMSO) for two hours on day *in vitro* (DIV) 10. In experiments to delineate signaling events downstream of 5-HT_2A_ receptor activation, neurons were treated with DOI (10 μM) for two hours in the presence of specific signaling pathway inhibitors, namely the mitogen activated protein kinase kinase (MAPKK) inhibitor U0126 (50 μM) and the CaM kinase II (CaMKII) inhibitor KN-62 (10 μM) (Tocris Bioscience, United Kingdom). The inhibitors were added to the cultures thirty minutes prior to DOI treatment, and were present throughout the duration of DOI exposure. Following treatments, cortical neurons were processed for immunofluorescence, RNA extraction for qPCR analysis, or western blot analysis.

### 2.3 Immunofluoresence

Immunofluorescence staining was performed as described previously (Fanibunda et al., 2019). In brief, cortical neurons were fixed in 4% paraformaldehyde, followed by blocking in 10% horse serum and incubated with primary antibodies, rabbit anti-pCREB (1:1000; Cell Signaling Technology, MA, USA) or goat anti-5-HT_2A_ receptor (1:500, Santa Cruz Biotechnologies, USA) along with the pan-neuronal marker, mouse anti-MAP2 (1:1000, Sigma-Aldrich, USA) overnight at 4°C. This was followed by incubation with secondary antibodies, Alexa 488 conjugated anti-goat (1:500; Molecular probes, CA, USA) or Alexa 488 conjugated anti-rabbit (1:500; Molecular probes, CA, USA) or biotinylated horse anti-mouse (1:500, Roche Applied Science, Switzerland) with subsequent incubation with streptavidin-conjugated Alexa 568 (1:500, Molecular probes, CA, USA) for two hours. Following secondary antibody incubation and serial washes, cortical neurons were mounted in Vectashield (Vector laboratories, CA, USA), and images were captured on the Zeiss LSM5 Exciter laser scanning microscope.

### 2.4 Western blot analysis

Rat cortical neurons were lysed in Laemmli sample buffer (2% SDS, 10% glycerol, 60 mM Tris-Cl, 0.01% bromophenol blue) and proteins were resolved via sodium dodecyl sulfate polyacrylamide gel electrophoresis (SDS-PAGE), followed by transfer onto polyvinylidene fluoride (PVDF, GE Healthcare, UK) membranes. Blots were blocked in 5% fat-free milk in 0.05% Tris Buffered Saline-Tween 20 (TBS-T) and incubated in primary antibodies in 0.05% TBS-T, overnight at 4°C. Primary antibodies included rabbit anti-pCREB (1:500; Cell Signaling Technology, MA, USA) and rabbit anti-CREB (1:1000, Cell Signaling Technology). Blots were washed 3–5 times and incubated with a 1:5000 dilution of horseradish peroxidase-conjugated goat anti-rabbit antibody (GE Healthcare, UK) for one hour. Protein-antibody complexes were detected on X-ray films following addition of Enhanced Chemiluminescence (ECL) substrate (GE Healthcare, UK). The relative density of the pCREB and CREB bands was quantitated using ImageJ software (NIH, USA), and was represented as a pCREB/CREB ratio.

### 2.5 qPCR analysis

RNA was isolated using Tri Reagent (Sigma-Aldrich, USA), according to the manufacturer’s protocols. 2 μg of RNA per sample was reverse transcribed using a complementary DNA (cDNA) synthesis kit (QuantiTect reverse transcription kit, Qiagen, Germany). Quantitative PCR (qPCR) was performed in a Mastercycler® ep realplex real-time PCR system (Eppendorf, Germany). cDNA was amplified using a SYBR Green kit (Applied Biosystems, CA, USA), with primers for the genes of interest (Supplementary Table 1). Hypoxanthine phosphoribosyl transferase (*Hprt*) was used as an endogenous housekeeping gene control for normalization. The relative expression levels between control and treated samples were computed by the comparative Ct method, as described previously (Schmittgen and Livak, 2008). Data are represented as fold change ± SEM as compared to control.

### 2.6 Chromatin immunoprecipitation

Chromatin immunoprecipitation (ChIP) was carried out as described previously (Desouza et al., 2011). Briefly, bilateral frontal neocortices were dissected and fixed to crosslink the DNA with the bound proteins. The tissue was placed in a pre-chilled Dounce homogenizer, sonicated and immunoprecipitated using a phosphorylated CREB (pCREB) antibody (1 μg; Cell Signaling Technology, MA, USA). Following reverse crosslinking and chromatin precipitation, qPCR analysis was performed for upstream sequences of the *Arc, BdnfI, Cebpb, cFos, Egr1* and *Egr2* promoters that contained putative CRE sites as predicted using AliBaba 2.1 (http://www.gene-regulation.com/pub/programs.html). For each sample the results were normalized to the input chromatin from the same sample. (Primer sequences used in ChIP experiments: Supplemental Table 2).

### 2.7 DOI-mediated head twitch behavior

Administration of DOI results in a stereotypical head twitch behavior, characterized by rapid radial movements of the head (Canal and Morgan, 2012; Halberstadt and Geyer, 2013). This behavior was videotaped in the home cage for a total duration of 20 minutes, commencing 20 minutes following administration of the DOI or vehicle (saline) treatment. The total number of head twitches in this time window were counted by an experimenter blind to the experimental treatment groups.

### 2.8. In situ hybridization

CREBαδ KO and litter-matched wild type (WT) mice were administered DOI and vehicle (Veh) resulting in four groups: WT + Veh, WT + DOI, CREBαδ KO + Veh, CREBαδ KO + DOI. Mice were anesthetized with sodium thiopentone and transcardially perfused with 4% paraformaldehyde (PFA). The brains were postfixed in 4% PFA overnight and cryoprotected in 30% sucrose in 4% PFA prior to being shipped to TIFR, India. Coronal sections of 30 μM thickness were cut on the freezing microtome (Leica Biosystems, Germany), fixed, blocked and acetylated. The floating sections were incubated for 20 hours at 60 °C in a hybridization buffer (50% formamide, 1xSSC, 25xDenhardt’s solution, 40 mM dithiothreitol, 150 μg/ml yeast tRNA, 10% dextran sulfate, 400 μg/ml salmon sperm DNA) containing ^35^S-UTP labeled antisense riboprobes for *Arc* mRNA at a concentration of 1×10^6^ cpm/300 μl. Antisense riboprobes to *Arc* mRNA were generated from a transcription-competent plasmid kindly provided by Dr. Oswald Steward (John Hopkins University, Baltimore). Following hybridization, all sections were washed in ribonuclease A (20 mg/ml; USB corporation, USA), followed by stringent washes in decreasing concentrations of SSC, mounted on slides, air dried and exposed to Hyper film β-max (GE Healthcare, USA) for seven days. Levels of Arc mRNA were quantified using Scion Image (Scion, USA) and calibrated using ^14^C standards to correct for nonlinearity. Equivalent areas of the somatosensory and prefrontal cortex were outlined and optical density measurements were determined (3–4 sections/animal).

### 2.9 Statistical analysis

Results were subjected to statistical analysis using Student’s unpaired *t*-test for experiments with two groups (GraphPad InStat) and one-way ANOVA (GraphPad, Prism 8) for experiments using signaling pathway inhibitors, followed by Tukey’s *post-hoc* test for group comparisons. For four group experiments statistical analysis was performed using two-way ANOVA (GraphPad, Prism 8). Tukey’s *post-hoc* test for group comparisons was applied when there was a significant two-way ANOVA interaction observed between the two variables of DOI treatment and CREBαδ KO genotype. Statistical significance was determined at *p* < 0.05.

## Results

### Acute treatment with the 5-HT_2A_ receptor agonist, DOI, regulates neuronal plasticity-associated gene expression via the MAP kinase and CaMKII signaling pathways and enhances phosphorylated CREB (pCREB) expression in vitro

The 5-HT_2A_ receptor agonist DOI, is a potent hallucinogen that is known to evoke an increase in the expression of several neuronal plasticity-associated genes (González-Maeso et al., 2003). Following ligand binding, the Gq-coupled 5-HT_2A_ receptor can differentially recruit multiple signaling pathways to bring about distinct signaling responses. We sought to address the contribution of the MAP kinase and CaM Kinase II (CaMKII) signaling pathways and the transcription factor CREB to the DOI-evoked induction of neuronal plasticity-associated gene expression. We stimulated primary rat cortical neurons *in vitro* with DOI, in the presence or absence of the CaMKII inhibitor, KN-62 or the MAPKK inhibitor, U0126 (Fig. 1A). We observed an upregulation of *Arc, Bdnf1, Cebpb* and *Egr2* mRNA levels following DOI treatment, which was inhibited by both the CaMKII inhibitor KN-62 and the MAPKK inhibitor U0126 (Fig. 1B) [One-way ANOVA: *Arc*: (F_(3,9)_ = 7.551, *p* = 0.008); *Bdnf1*: (F_(3,11)_ = 6.436, *p* = 0.009); *Cebpb*: (F_(3,8)_ = 5.293, *p* = 0.027), *Egr2*: (F_(3,10)_ = 7.685, *p* = 0.006)]. In contrast, the DOI-mediated upregulation of *cFos* mRNA levels was prevented by only the KN-62 inhibitor, while the increase in the *Egr1* transcript levels was abrogated exclusively by the U0126 inhibitor (Fig. 1B) [One-way ANOVA: *cFos*: (F_(3,11)_ = 5.563, *p* = 0.014), *Egr1*: (F_(3,11)_ = 14.02, *p* = 0.0004)]. This indicates a differential involvement of the MAP kinase and the CaMKII pathway in the 5-HT_2A_ receptor-mediated transcriptional regulation of specific neuronal plasticity-associated genes (Fig. 1C).

**Figure 1.**
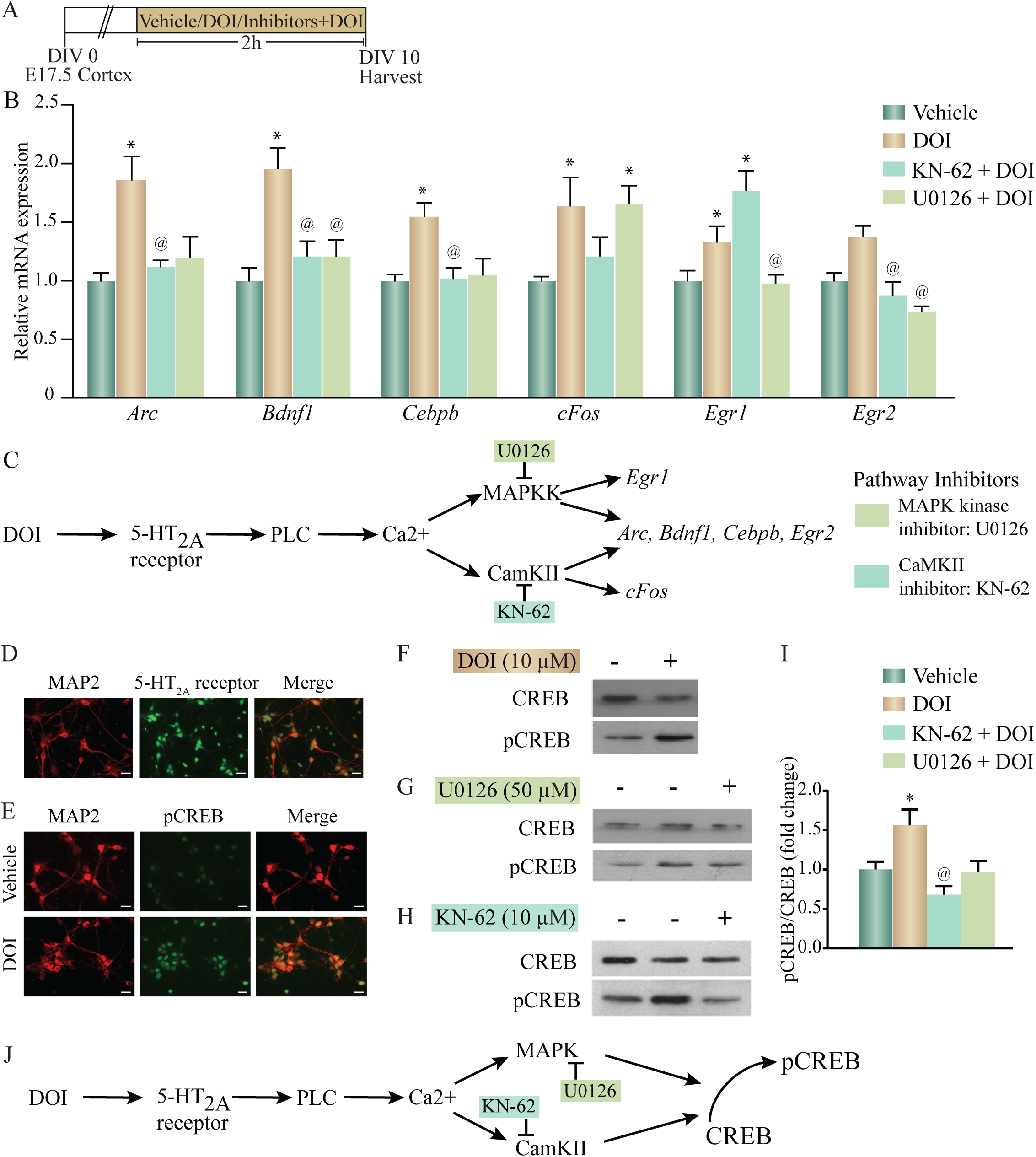
Acute treatment with the 5-HT_2A_ receptor agonist, DOI regulates neuronal plasticity-associated gene expression via the MAP kinase and CaMKII signaling pathways and enhances phosphorylated CREB (pCREB) expression in vitro. (A) Shown is a schematic of the treatment paradigm for cortical neurons derived from E17.5 rat embryos, allowed to differentiate till day *in vitro* (DIV) 10, following which neurons were treated with vehicle (DMSO) or the 5-HT_2A_ receptor agonist, DOI (10 μM), in the presence or absence of CaMKII and MAP kinase signaling pathway inhibitors (CaMKII inhibitor: KN-62; MAPKK inhibitor: U0126). (B) Shown is the relative mRNA expression for plasticity-associated genes following DOI treatment in the presence or absence of MAP kinase and CaMKII signaling pathway inhibitors, represented as fold change of vehicle ± SEM. (Representative results from n = 3 - 5 wells per treatment group/N = 3, \**p* < 0.05 as compared to vehicle, ^@^*p* < 0.05 as compared to DOI, one-way ANOVA, Tukey’s *post-hoc* test). (C) Shown is a schematic summarizing the putative signaling pathways that may contribute to DOI-induced gene expression. The CaMKII inhibitor, KN-62 and the MAPKK inhibitor, U0126 inhibit the CaMKII and MAP kinase signaling pathways respectively. The DOI-mediated upregulation of *Arc, Bdnf1, Cebpb* and *Egr2* mRNA levels was blocked by both the MAPKK and CaMKII inhibitors, whereas the increase in *cFos* mRNA was blocked by the CaMKII, not the MAPKK, inhibitor and the upregulation of *Egr1* mRNA was blocked by the MAPKK, not the CaMKII, inhibitor. (D) Shown are representative immunofluorescence images of rat cortical neurons *in vitro* with double staining for the neuronal marker MAP2 (red) and 5-HT_2A_ receptor (green). Scale bar: 30 µm. Magnification: 20X. (E) Shown are representative immunofluorescence images of rat cortical neurons with double staining for pCREB (green) and the neuronal marker MAP2 (red) - upper panel: Vehicle; lower panel: DOI. Scale bar: 30 µm. Magnification: 20X. (F-I) Shown are representative immunoblots for pCREB and CREB protein levels in rat cortical neurons treated with DOI (F) or with DOI in the presence or absence of the MAPKK inhibitor U0126 (G) or the CaMKII inhibitor KN-62 (H). (I) Quantitative densitometric analysis of pCREB/CREB levels in rat cortical neurons treated with DOI in the presence or absence of U0126 or KN-62. Results are expressed as fold change of vehicle ± SEM. (Representative results from n = 3 - 5 wells per treatment group/N = 3, \**p* < 0.05 as compared to vehicle, ^@^*p* < 0.05 as compared to DOI, one-way ANOVA, Tukey’s *post-hoc* test). (J) Shown is a schematic depicting the putative pathway via which pCREB levels are enhanced following DOI administration, indicative of a role for the MAP kinase and CaMKII signaling pathways.

We next sought to address the possible role of the transcription factor CREB in mediating the signaling events evoked by DOI, downstream of the 5-HT_2A_ receptor. The DOI regulated genes, *Arc, Bdnf1, Cebpb, cFos, Egr1* and *Egr2* were found to contain putative cAMP response element (CRE) (TGACG/CGTCA/TGACGTCA) sites in the upstream promoter regions, suggestive of the possibility of a CREB-dependent transcriptional regulation. We first confirmed 5-HT_2A_ receptor expression in cortical neurons, by immunofluorescence staining (Fig. 1D). Immunostaining with the phosphorylated CREB (pCREB) antibody demonstrated an increase in pCREB immunofluorescence intensity in DOI-treated rat cortical neurons as compared to vehicle-treated control neurons (Fig. 1E). Immunoblotting to detect pCREB levels, also demonstrated a robust increase in pCREB/CREB levels in DOI-treated cortical neurons (Fig. 1F), further corroborating that DOI treatment enhances pCREB levels. Further, the MAPKK inhibitor (Fig. 1G) and the KN-62 inhibitor (Fig. 1H), both prevented the DOI-evoked increase in pCREB/CREB levels, demonstrating a role for the MAP kinase pathway and CaMKII pathway in mediating the DOI-evoked increase in pCREB levels (Fig. 1I) (pCREB/CREB: F_(3,17)_ = 5.561, *p* = 0.008). This data collectively suggests that DOI, a hallucinogenic ligand of the Gq-coupled 5-HT_2A_ receptor, recruits MAP kinase and CaMKII signaling to enhance the phosphorylation of the transcription factor CREB in rat cortical neurons (Fig.1J).

### Acute treatment with DOI enhances both the expression of putative CRE-containing plasticity-associated genes and the enrichment of pCREB within the promoter regions of specific plasticity-associated genes in the neocortex of adult rats

Given that we observed that the 5-HT_2A_ receptor agonist, DOI, induced a robust upregulation of neuronal plasticity-associated genes in rat cortical neurons in culture, we next examined whether DOI administration to Sprague-Dawley male rats could evoke similar alterations in neuronal plasticity-associated genes in the neocortex. We systemically administered DOI (8 mg/kg) and evaluated gene expression in the neocortex at a two hour time-point post treatment (Fig. 2A). DOI administration evoked a robust increase in the gene expression of *Arc, Atf3, Atf4, Bdnf1, Cebpb, Cebpd, Egr1, Egr2, Egr3, Egr4, cFos, JunB* and *Nfkbia* in the rat neocortex (Fig. 2B).

**Figure 2:**
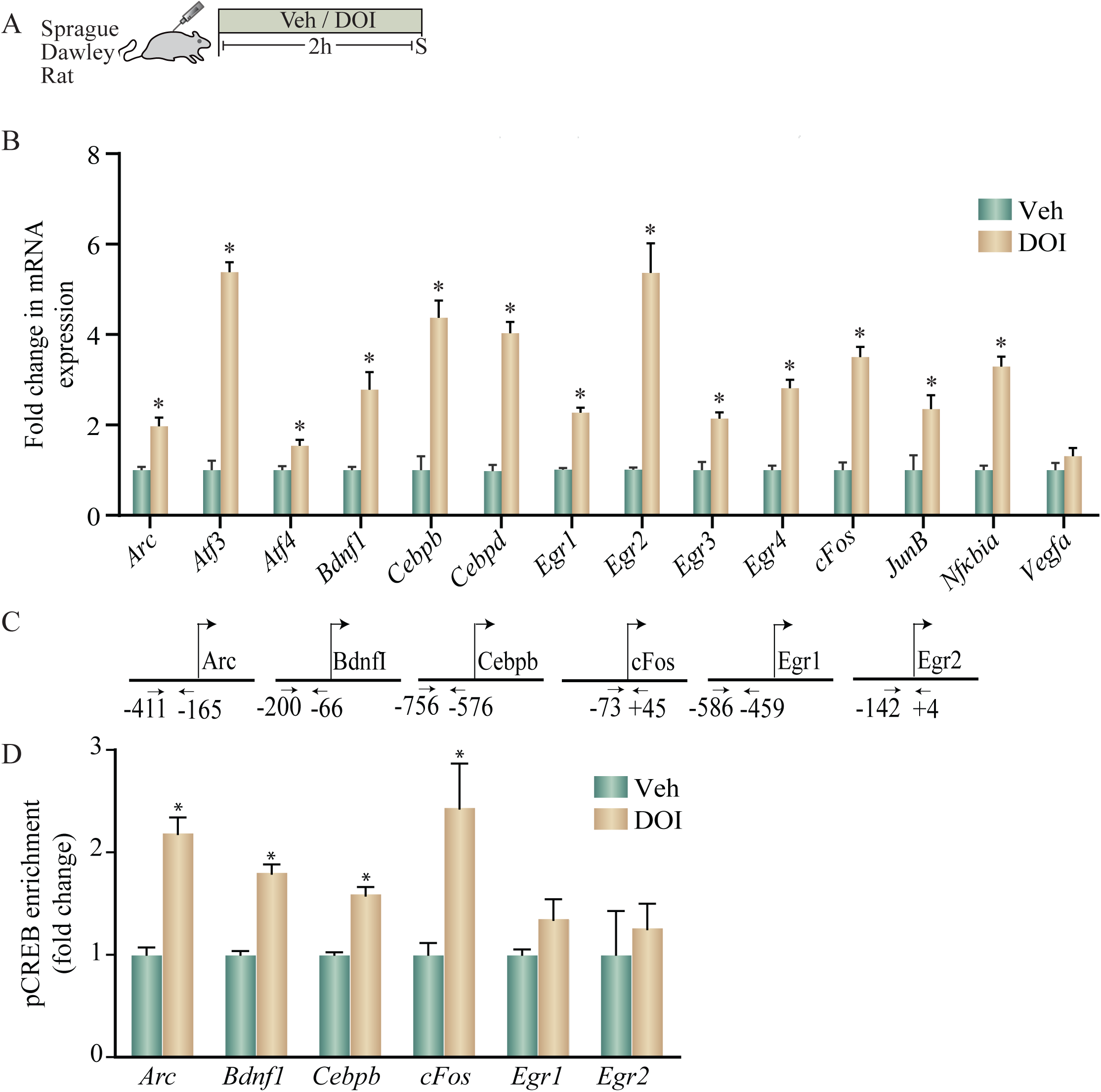
Acute treatment with DOI enhances both the expression of putative CRE-containing plasticity-associated genes and the enrichment of pCREB within the promoter regions of specific plasticity-associated genes in the neocortex of adult rats. (A) Shown is a schematic of the treatment paradigm wherein adult Sprague-Dawley rats were injected with vehicle or DOI (8 mg/kg) and were sacrificed two hours following treatment. (B) The bar graph indicates the fold change in mRNA expression of specific plasticity-associated genes in the neocortex of vehicle and DOI-treated rats represented as fold change of vehicle ± SEM (n = 4 - 6 per treatment group, \**p* < 0.05 as compared to vehicle, unpaired Students *t*-test). (C) Shown are the chromatin immunoprecipitation (ChIP) PCR amplicons with primer locations spanning putative CRE sequences in the upstream gene regulatory sequences for *Arc, Bdnf1, Cebpb, cFos, Egr1* and *Egr2*. (E) Shown is a bar graph for pCREB enrichment at the *Arc, Bdnf1, Cebpb, cFos, Egr1* and *Egr2* promoters based on ChIP analysis performed on tissue derived from the neocortex of vehicle and DOI treated adult rats. Results are expressed as the fold change of vehicle ± SEM. (n = 7 - 10 animals per treatment group, \**p* < 0.05 as compared to vehicle, unpaired Students *t*-test).

We next examined if a subset of the genes upregulated by DOI treatment, that are known to contain putative CRE binding sites in their promoter regions based on *in silico* analysis, also exhibit enrichment for pCREB within their promoters (Fig. 2C). Enhanced expression of *Arc, Bdnf1, Cebpb* and *cFos* in the neocortex of DOI-treated animals was accompanied by a significant enrichment of pCREB at the promoters of *Arc, Bdnf1, Cebpb* and *cFos* (Fig. 2D). In contrast, the enhanced gene expression of *Egr1* and *Egr2* was not associated with any significant change in pCREB enrichment at putative CRE sites within their promoter regions (Fig. 2D). Taken together, these results indicate that *in vivo* administration of the 5-HT_2A_ receptor agonist DOI, evokes a robust upregulation of several neuronal plasticity-associated genes in the neocortex, a subset of which exhibit significant pCREB enrichment in their promoter regions.

### DOI-mediated regulation of plasticity-associated gene expression in cortical brain regions is perturbed in CREBαδ knockout (CREBαδ KO) mice

Given the evidence both *in vitro* and *in vivo*, that the 5-HT_2A_ receptor agonist, DOI induces (1) a robust increase in pCREB levels in rat cortical neurons, (2) a significant enrichment of pCREB at the promoters of specific plasticity-associated genes that are robustly enhanced by DOI treatment, we next sought to address the contribution of CREB to the regulation of these transcripts. We used hypomorphic CREBαδ KO mice (Blendy et al., 1996) that are reported to have a greater than 90% reduction in CREB binding activity to consensus CRE target sites (Walters and Blendy, 2001). We treated WT and CREBαδ KO mice with vehicle or DOI (8 mg/kg), and assessed transcript expression of specific neuronal plasticity-associated genes at a two hour time-point post treatment (Fig. 3A). To rule out the possibility that CREB may indirectly regulate 5-HT_2A_ or 5-HT_2C_ receptors, we first assessed whether the baseline expression of these receptors was altered in CREBαδ KO mice as compared to their WT controls, and observed no change in 5-HT_2A_ or 5-HT_2C_ receptor mRNA levels within the neocortex (Fig. 3B). Further to assess whether CREB deficient mice exhibit any change in 5-HT_2A_ receptor evoked behavioral responses, we evaluated a stereotypical head twitch response (HTR) behavior evoked by the 5-HT_2A_ receptor agonist DOI (Canal and Morgan, 2012; Halberstadt and Geyer, 2013). Behavioral analysis to quantify the number of HTR events in WT and CREBαδ KO mice, indicated that the number of HTR responses evoked by DOI were unaltered in CREBαδ KO mice (Fig. 3C). While we noted no significant two-way ANOVA interaction of DOI and CREBαδ KO genotype, we observed a significant main effect of DOI (F_(1,12)_ = 58.64, *p* < 0.0001). This indicates that the loss of CREBαδ isoforms does not alter either the expression of the 5-HT_2A_ receptor, or the behavioral head twitch responses evoked by the 5-HT_2A_ receptor agonist DOI, that are critically dependent on the cortical 5-HT_2A_ receptor.

**Figure 3:**
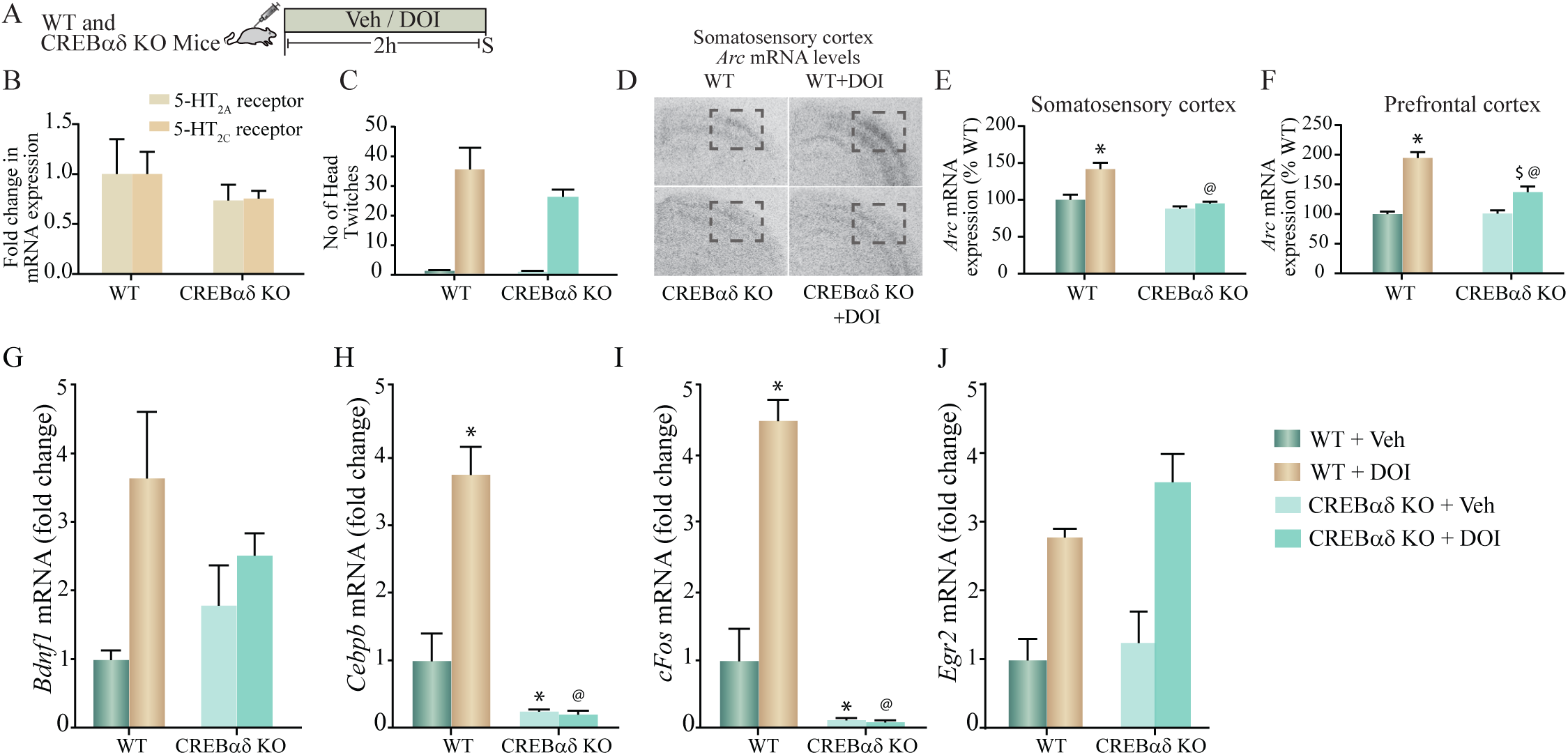
DOI-mediated regulation of plasticity-associated gene expression in cortical brain regions is perturbed in CREBαδ knockout (CREBαδ KO) mice. (A) Shown is a schematic of the acute treatment paradigm for wild type (WT) and CREBαδ KO mice, with vehicle (saline) or DOI (8 mg/kg) followed by sacrifice two hours after treatment. (B) The bar graph depicts quantitative qPCR analysis for 5-HT_2A_ and 5-HT_2C_ receptors in the neocortex of WT and CREBαδ KO mice represented as fold change of WT ± SEM. (n = 3 - 4 animals per treatment group). (C) The bar graph depicts the quantitation of head-twitch responses evoked in response to acute treatment with the 5-HT_2A_ receptor agonist, DOI or vehicle in both WT and CREBαδ KO mice (n = 4 animals per treatment group) (D) Shown are representative autoradiographs for *Arc* mRNA expression in the neocortex from WT, WT + DOI, CREBαδ KO and CREBαδ KO + DOI mice, with the outline inset indicating the somatosensory region. Shown are bar graphs for the quantitative densitometric analysis of levels of *Arc* mRNA expression in the somatosensory (E) and prefrontal (F) cortex following DOI or vehicle treatment to WT and CREBαδ KO mice. Results are represented as percentage of WT and are mean ± SEM (n= 3 - 4 animals per group, \**p* < 0.05 as compared to WT + Veh mice, ^$^*p* < 0.05 as compared to CREBαδ KO + Veh mice, ^@^*p* < 0.05 as compared to WT + DOI mice, two-way ANOVA, Tukey’s *post-hoc* test). (G-J) Bar graphs depict quantitation of qPCR analysis for mRNA expression of *Bdnf1* (G), *Cebpb* (H), *cFos* (I) and *Egr2* (J), following acute DOI or vehicle treatment to WT and CREBαδ KO mice. (n = 3 - 4 animals per group, \**p* < 0.05 as compared to WT mice, ^$^*p* < 0.05 as compared to CREBαδ KO mice, ^@^*p* < 0.05 as compared to WT+DOI mice, two-way ANOVA, Tukey’s *post-hoc* test).

We next performed *in situ* hybridization and qPCR analysis to address whether the regulation of specific neuronal plasticity-associated genes that exhibit pCREB enrichment at their promoters following DOI treatment, were altered in the neocortex of CREBαδ KO mice. Radioactive *in situ* hybridization analysis indicated that the DOI-evoked robust induction of *Arc* mRNA levels both in the somatosensory cortex and in the prefrontal cortex was significantly attenuated in CREBαδ KO mice (Fig. 3D). Two-way ANOVA analysis for *Arc* mRNA levels in the somatosensory cortex (Fig. 3E) indicated a significant DOI by CREBαδ KO genotype interaction (F_(1,12)_ = 8.39, *p* = 0.013), as well as significant main effects of DOI (F_(1,12)_ = 16.81, *p* = 0.002) and CREBαδ KO genotype (F_(1,12_) = 24.39, *p* = 0.0003). The robust DOI-evoked upregulation of *Arc* mRNA in the somatosensory cortex of WT control mice was completely lost in the CREBαδ KO mice. Two-way ANOVA analysis for *Arc* mRNA levels in the prefrontal cortex (Fig. 3F) indicated a significant DOI by CREBαδ KO genotype interaction (F_(1,11)_ = 15.73, *p* = 0.002), as well as significant main effects of DOI (F_(1,11)_ = 78.83, *p* = 0.0001) and CREBαδ KO genotype (F_(1,11)_ = 14.75, *p* = 0.003). The robust DOI-evoked upregulation of *Arc* mRNA in the prefrontal cortex of WT control mice was significantly attenuated in the CREBαδ KO mice. No change was observed in the basal expression of *Arc* mRNA in either the somatosensory or prefrontal of CREBαδ KO mice, which did not differ from vehicle-treated WT controls.

qPCR analysis was carried out to assess whether the DOI-evoked upregulation of *Bdnf1, Cebpb, cFos* and *Egr2* mRNA expression was altered in the neocortex of CREBαδ KO mice following acute DOI administration (Fig. 3G-J). Two-way ANOVA analysis for *Bdnf1* mRNA levels in the neocortex indicated no significant DOI by CREBαδ KO genotype interaction (Fig. 3G), however we did observe a significant main effect of DOI (F_(1,10)_ = 10.19, *p* = 0.01) and no main effect of CREBαδ KO genotype. Two-way ANOVA analysis for *Cebpb* mRNA levels in the neocortex (Fig. 3H) indicated a significant DOI by CREBαδ KO genotype interaction (F_(1,12)_ = 6.413, *p* = 0.026), as well as a trend towards a main effect of DOI (F_(1,12)_ = 3.388, *p* = 0.09) and a significant main effect of CREBαδ KO genotype (F_(1,12)_ = 50.49, *p* = 0.0001). Tukey’s *post-hoc* group comparisons indicated that the CREBαδ KO mice exhibited a significant baseline decrease (*p* = 0.03) in *Cebpb* mRNA levels in the neocortex. Further, while DOI-evoked a robust and significant upregulation of *Cebpb* mRNA levels in WT animals (*p* = 0.04), this was completely lost in the CREBαδ KO mice. *Cebpb* mRNA levels in the DOI-treated WT cohort differed significantly from the DOI-treated CREBαδ KO mice (*p* < 0.0001). Two-way ANOVA analysis for *cFos* mRNA levels in the neocortex (Fig. 3I) indicated a significant DOI by CREBαδ KO genotype interaction (F_(1,12)_ = 8.966, *p* = 0.011), as well as a trend towards a main effect of DOI (F_(1,12)_ = 4.493, *p* = 0.056) and a significant main effect of CREBαδ KO genotype (F_(1,12)_ = 102.5, *p* < 0.0001). Tukey’s *post-hoc* group comparisons indicated that the CREBαδ KO mice exhibited a significant baseline reduction (*p* = 0.001) in expression of *cFos* mRNA in the neocortex, and while DOI-evoked a robust and significant upregulation of *cFos* mRNA levels in WT animals (*p* = 0.02) this was completely lost in the CREBαδ KO mice. Neocortical *c-Fos* mRNA levels differed significantly between DOI-treated WT mice and the DOI-treated CREBαδ KO cohort (*p* < 0.0001). In contrast, we observed no significant DOI by CREBαδ KO genotype interaction for *Egr2* mRNA levels within the neocortex (Fig. 3J). Further, while we did note a significant main effect of DOI for *Egr2* expression (F_(1,11)_ = 10.19, *p* = 0.003) we observed no main effect of CREBαδ KO genotype. Collectively, these observations indicate that CREB contributes to the acute DOI-evoked upregulation of *Arc, Cebpb* and *cFos* mRNA within the neocortex, but not to the DOI-induced neocortical increase in *Bdnf1* and *Egr2* mRNA expression. Furthermore, baseline expression of *Cebpb* and *cFos* mRNA, but not *Arc* mRNA is also significantly attenuated in the CREBαδ KO mice. These results demonstrate that the transcription factor CREB contributes to the regulation of a subset of neuronal plasticity-associated genes that are regulated by the hallucinogenic 5-HT_2A_ receptor agonist, DOI in the neocortex.

## 4. Discussion

Here, we show that DOI, a hallucinogenic ligand of the Gq-coupled 5-HT_2A_ receptor, recruits the nuclear transcription factor CREB to influence the expression of a specific subset of neuronal plasticity-associated genes in the rodent neocortex. Stimulation of the 5-HT_2A_ receptor in rat cortical neurons recruits the MAP kinase and CaMKII signaling pathways to rapidly increase the phosphorylation of CREB. 5-HT_2A_ receptor stimulation by DOI, results in the upregulation of *Arc, Bdnf1, Cebpb* and *Egr2* mRNA in cortical neurons, with a role for both the MAP kinase and CaMKII signaling pathways. We also noted that while the regulation of *Egr1* is dependent on the MAP kinase pathway, the regulation of *cFos* expression by DOI is dependent on the CaMKII pathway. Further, we show recruitment of pCREB at CRE binding sites within the promoters of a subset of DOI-regulated target genes (*Arc, Bdnf1, Cebpb, cFos*) in the neocortex. In contrast, there are also specific DOI-regulated plasticity-associated genes (*Egr1, Egr2)* wherein we did not observe pCREB enrichment at putative CRE sites in their promoters. These findings collectively indicate that 5-HT_2A_ receptor stimulation enhances pCREB enrichment at the promoters of a subset of DOI regulated target genes, suggestive of an upregulation of neuronal plasticity-associated genes in both a CREB-dependent and independent manner. This is further supported by evidence that the DOI-evoked transcriptional upregulation of specific neuronal plasticity-associated genes (*Arc, cFos* and *Cebpb*) is attenuated in the neocortex of CREBαδ KO mice (Fig. 4).

**Figure 4:**
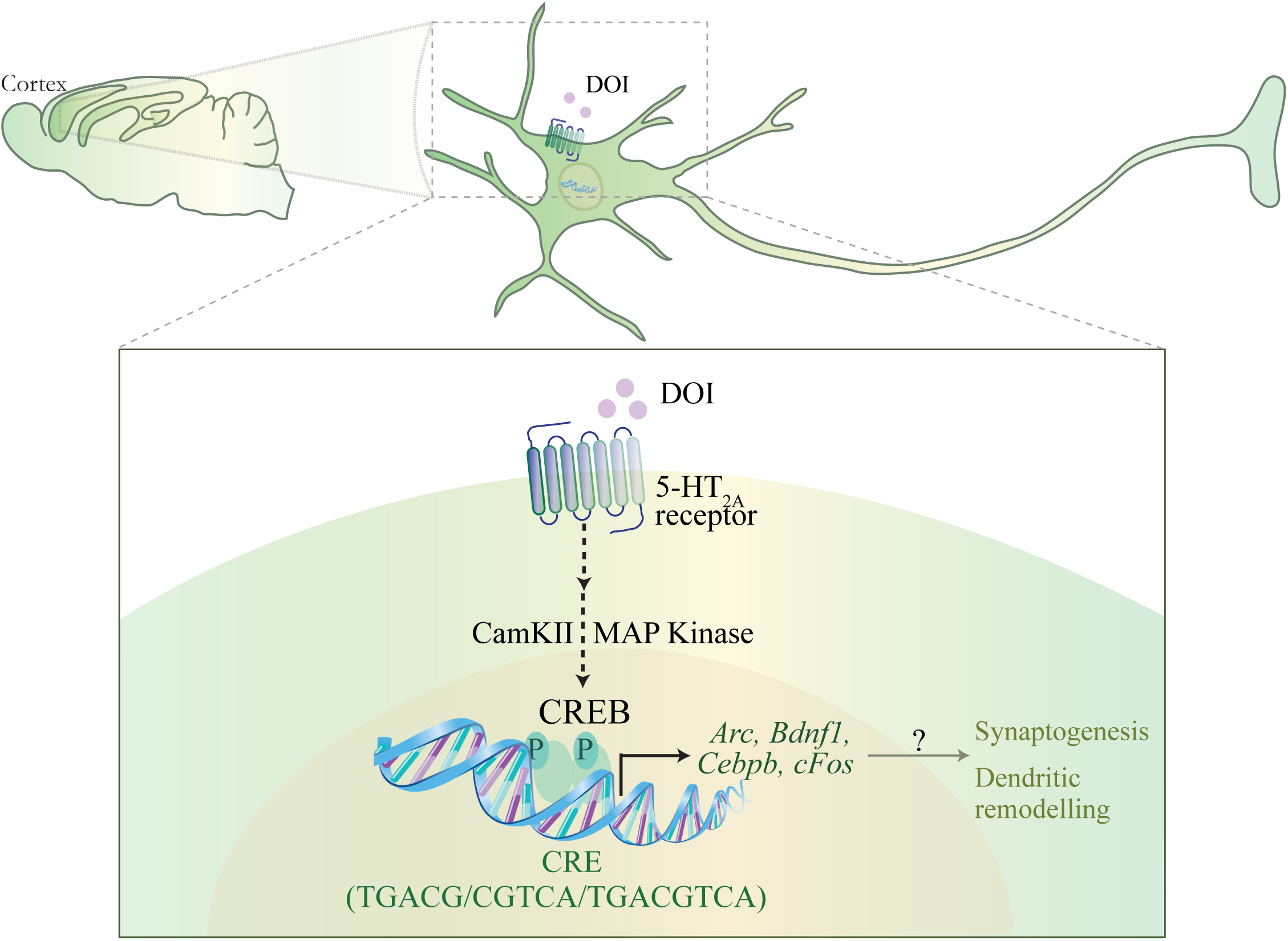
Schematic depicting the putative mechanism for the CREB-dependent regulation of neuronal plasticity-associated gene expression by the hallucinogenic 5-HT_2A_ receptor agonist DOI. DOI, a hallucinogenic agonist of the 5-HT_2A_ receptor is known to evoke a specific transcriptome signature within the neocortex, including the upregulation of the expression of several plasticity-associated genes. The schematic indicates a putative mechanism through which DOI-mediated stimulation of the 5-HT_2A_ receptor, results in the recruitment of the MAP kinase and CaMKII signaling pathways, which would further result in the phosphorylation of the transcription factor CREB, thus facilitating the CREB-dependent transcription of plasticity-associated genes, *Arc, Bdnf1, Cebpb, cFos*. This raises the intriguing possibility that CREB-dependent regulation of gene expression could contribute to the effects of the hallucinogenic 5-HT_2A_ receptor agonist DOI, on neuronal plasticity, synaptogenesis and cell survival.

Prior literature indicates that ligands at the 5-HT_2A_ receptor can differentially recruit unique transcriptome fingerprints in the neocortex (González-Maeso et al., 2003), with the hallucinogenic compounds DOI and LSD enhancing the expression of specific transcripts, including *Egr1* and *Egr2* (González-Maeso et al., 2003, 2007). Despite the knowledge that hallucinogenic agonists of the 5-HT_2A_ receptor, such as DOI, induce the expression of multiple neuronal plasticity-associated genes, the contribution of specific signaling pathways and transcription factors to the 5-HT_2A_ receptor regulated cortical gene expression remains poorly elucidated. DOI-mediated 5-HT_2A_ receptor stimulation recruits the phospholipase C (PLC)-PKC-MAP kinase cascade and the PLC-CaMKII pathways that are known to also target the transcription factor CREB (Banerjee and Vaidya, 2020; Sharp and Barnes, 2020). The beta-arrestin-receptor complex also acts as a scaffold to activate the Raf-MEK-MAP kinase cascade (Schmid et al., 2008; McCorvy and Roth, 2015). Following 5-HT_2A_ receptor activation by DOI, we observe that both the MAP kinase and CaMKII pathways, contribute to the DOI-evoked transcriptional increase of *Arc, Bdnf1, Cebpb* and *Egr2* expression in rat cortical neurons, whilst the CaMKII pathway and the MAP kinase pathway contribute to the DOI-mediated regulation of *cFos* and *Egr1* expression respectively. This highlights the recruitment of distinct signaling pathways by the 5-HT_2A_ receptor agonist, DOI (Banerjee and Vaidya, 2020), several of which converge on the transcription factor CREB (Shaywitz and Greenberg, 1999), placing it as a potential key ‘hub’ regulator of the transcriptional changes that arise in response to DOI treatment. Interestingly, many of the genes we find to be robustly enhanced in expression following DOI treatment in cortical neurons *Arc, Bdnf1, Cebpb, cFos, Egr1* and *Egr2*, are reported to contain putative CRE sites (TGACG/ CGTCA/ TGACGTCA) (Sheng et al., 1990; Niehof et al., 1997; Ahn et al., 1998; Walters and Blendy, 2001; Fukuchi et al., 2005; Bilbao et al., 2014; Duclot and Kabbaj, 2017; Ly et al., 2020). Our *in vitro* studies clearly indicate a robust induction in pCREB levels at the Ser133 site, with phosphorylation via the MAP kinase and CaMKII pathways likely contributing to this induction. Our experiments motivate further characterization of the target sites of CREB phosphorylation (Kornhauser et al., 2002), as certain phosphorylation signatures like Ser142 (Sun et al., 1994) can also serve to evoke an inhibitory effect on CREB mediated transcription.

A previous microarray study from our lab has reported several genes to be upregulated by DOI in the rodent neocortex (Benekareddy et al., 2010). We performed *in silico* analysis on the published array results, and noted that several of the genes upregulated by DOI have CRE elements in their upstream regulatory promoter regions, suggestive of a putative role for CREB in the regulation of multiple DOI-evoked transcripts. DOI-evoked a robust increase in the transcript expression of several CRE-containing genes, namely *Arc, Atf3, Atf4, Bdnf1, Cebpb, Cebpd, Egr1, Egr2, Egr3, Egr4, cFos, JunB* and *Nfkbia* in the rat neocortex. Interestingly, we observed pCREB enrichment at the promoters of specific DOI-regulated genes. The expression of *Arc, Bdnf1, Cebpb* and *cFos* genes but not *Egr1* and *Egr2* expression was accompanied by a significant enrichment of pCREB at their promoters in the neocortex of DOI-treated animals. In this regard, it is interesting to note that the genes, *Egr1* and *Egr2* which are part of the reported DOI-evoked hallucinogenic fingerprint (González-Maeso et al., 2003) do not appear to show pCREB enrichment at their promoters. While we have restricted our analysis of pCREB recruitment to CRE sites close to the promoter region, we cannot preclude the possibility of CREB binding at remote CRE sites either in distal enhancer regions, or within introns as we have not scanned pCREB enrichment at these loci. It is also important to keep in mind that the differential regulation of target genes by CREB may further be influenced by CRE sequence composition, location of the CRE and distance from the transcription start site (Mayr and Montminy, 2001; Altarejos and Montminy, 2011; Davis et al., 2020). Thus, the presence of a CRE site does not necessarily predict the recruitment of pCREB, and other transcription factors besides CREB are also likely to be recruited by DOI and by neuronal activity-dependent mechanisms, to evoke transcriptional increase of target genes. Our findings provide impetus for a broader genome-wide analysis of pCREB enrichment to get a sense of the span of pCREB-mediated regulation of transcription by DOI, and an understanding of recruitment of pCREB at both canonical and non-canonical CREs.

It is also of interest to note that the target genes regulated by DOI are known to exhibit substantial signaling crosstalk, and could also exert further feedback effects on CREB-dependent transcriptional regulation (Wu et al., 2001; Deisseroth and Tsien, 2002; Wiegert and Bading, 2011; Belgacem and Borodinsky, 2017). *Bdnf*, a DOI-regulated target gene, also contributes to the transcriptional regulation of *Arc* mRNA expression (Bramham et al., 2008; Benekareddy et al., 2012), and could further via regulation of the MAP kinase cascade impinge on pCREB mediated gene regulation (Finkbeiner et al., 1997). This raises the intriguing possibility that reciprocal interactions between BDNF and CREB could serve to amplify the regulation of several neuronal plasticity-associated genes (Finkbeiner et al., 1997; Shieh and Ghosh, 1999; Nair and Vaidya, 2006). These include the transcriptional regulation of *Arc* and *c-Fos* which are reported to play a significant role in coupling experience-dependent transcriptional regulation to synaptic plasticity (Duman RS, 2005; Bramham et al., 2008; Flavell and Greenberg, 2008; Shepherd and Bear, 2011; Minatohara et al., 2015). Fos may also function at enhancer elements to coordinate global activity-dependent gene transcription (Malik et al., 2014; Joo et al., 2016). This then suggests that stimulation of the 5-HT_2A_ receptor, sets into play a coordinated transcriptional program that involves crosstalk of diverse signaling cascades and transcription factors that drive the expression of several neuronal plasticity-associated genes (*Arc, Bdnf1, cFos*), with CREB being amongst the key hub transcriptional factors that contributes to an important component of the DOI-evoked gene regulation pattern.

We have capitalized on the use of the CREBαδ KO mouse line, which is deficient for the α and δ CREB isoforms leading to a robust reduction in CREB binding to consensus CRE target sites (Walters and Blendy, 2001), to evaluate the contribution of CREB to the effects of DOI on neuronal plasticity-associated gene expression. It is important to note that the CREBαδ KO mice do not exhibit alterations in either the baseline expression of the 5-HT_2A_ receptor, or the stereotypical head twitch response (HTR) behavior evoked by the 5-HT_2A_ receptor agonist DOI, that is known to be dependent on the cortical 5-HT_2A_ receptor. This would suggest that alterations in DOI-evoked gene expression in CREBαδ KO mice are unlikely to arise due to a change at the level of 5-HT_2A_ receptor expression and coupling given that the CREBαδ KO mice exhibit a robust DOI-evoked HTR response no different from WT controls and show no change in 5-HT_2A_ receptor expression. Interestingly, the DOI-evoked induction in cortical *Arc* mRNA expression was lost in CREBαδ KO mice. Similar to this observation, the DOI-evoked increase in *Cebpb* and *cFos* mRNA levels were also abolished in the neocortex of CREBαδ KO mice. Further, in CREB deficient mice the baseline expression of *Cebpb* and *cFos*, but not *Arc* mRNA was also reduced, thus supporting a role for CREB in regulation of basal expression of *Cebpb* and *cFos*. The DOI-evoked induction in *Egr2* mRNA was not altered in the neocortex of CREBαδ KO mice, which is consistent with the evidence that pCREB was not found to be enriched at the *Egr2* promoter following DOI treatment. The DOI-evoked increase in *Bdnf1* transcript variant levels was also not significantly attenuated in CREBαδ KO mice, and further we did not observe a change in basal *Bdnf1* transcript expression either. We have focused on *Bdnf1* which has been reported to contain CRE elements at its promoter, and exhibit CREB-mediated regulation in other contexts (Tabuchi et al., 2002; Esvald et al., 2020). *BdnfIII* and *BdnfIV* promoters are also known to contain CRE elements, as is the *Bdnf* coding exon, and it will be important to systematically address the contribution of CREB to the DOI-mediated regulation of multiple *Bdnf* transcript variants (Tao et al., 1998; Esvald et al., 2020). Indeed, prior reports clearly indicate several conditions in which CREB-dependent regulation of *Bdnf* gene expression contributes to neuroplasticity, in particular in the context of neuronal activity-dependent transcriptional coupling that drives structural and synaptic plasticity (Shieh and Ghosh, 1999; Vogt et al., 2014; Yan et al., 2016; Rafa-Zabłocka et al., 2018). While the CREB deficient mice provide a valuable tool to address the contribution of CREB to the DOI-mediated regulation of plasticity-associated gene expression, they come with a caveat of a constitutive, developmental onset loss-of-function of CREB which could in turn disrupt several key signaling pathways. Further experiments are warranted to systematically evaluate the contribution of CREB to the effects of DOI on neuronal plasticity-associated genes using approaches that allow for a more targeted strategy of adult onset, neuronal circuit-specific loss of function of CREB.

Our findings highlight a key role for CREB in contributing to the DOI-mediated regulation of specific neuronal plasticity-associated genes in the neocortex. This raises the intriguing possibility that similar to slow-onset, pharmacological antidepressants that recruit CREB to drive transcriptional changes in neurotrophic and plasticity-associated genes (Nibuya et al., 1996; Duman RS, Nibuya M, 1997; Thome et al., 2000; Chen et al., 2001), serotonergic psychedelics that target the 5-HT_2A_ receptor may also recruit CREB, albeit more rapidly, to drive a plasticity-associated transcriptional program. These observations encourage further investigation into the role of CREB in regulating the transcription of plasticity-associated genes evoked by hallucinogenic 5-HT_2A_ receptor agonists, thus creating a conducive milieu for the psychoplastogenic actions of serotonergic psychedelics on dendritic plasticity and synaptogenesis in the neocortex.

## 6. Author Contributions

LD, MB, SF, MF, TG - performed experiments and analyzed the data; JB, VV - designed experiments and analyzed data; SF, VV - wrote the manuscript.

## 8. Acknowledgements

We thank Dr. Shital Suryavanshi and the animal house staff at the Tata Institute of Fundamental Research (TIFR), Mumbai for technical assistance.

## 10. Supplementary material

### List of Primers

**Supplementary Table 1:**
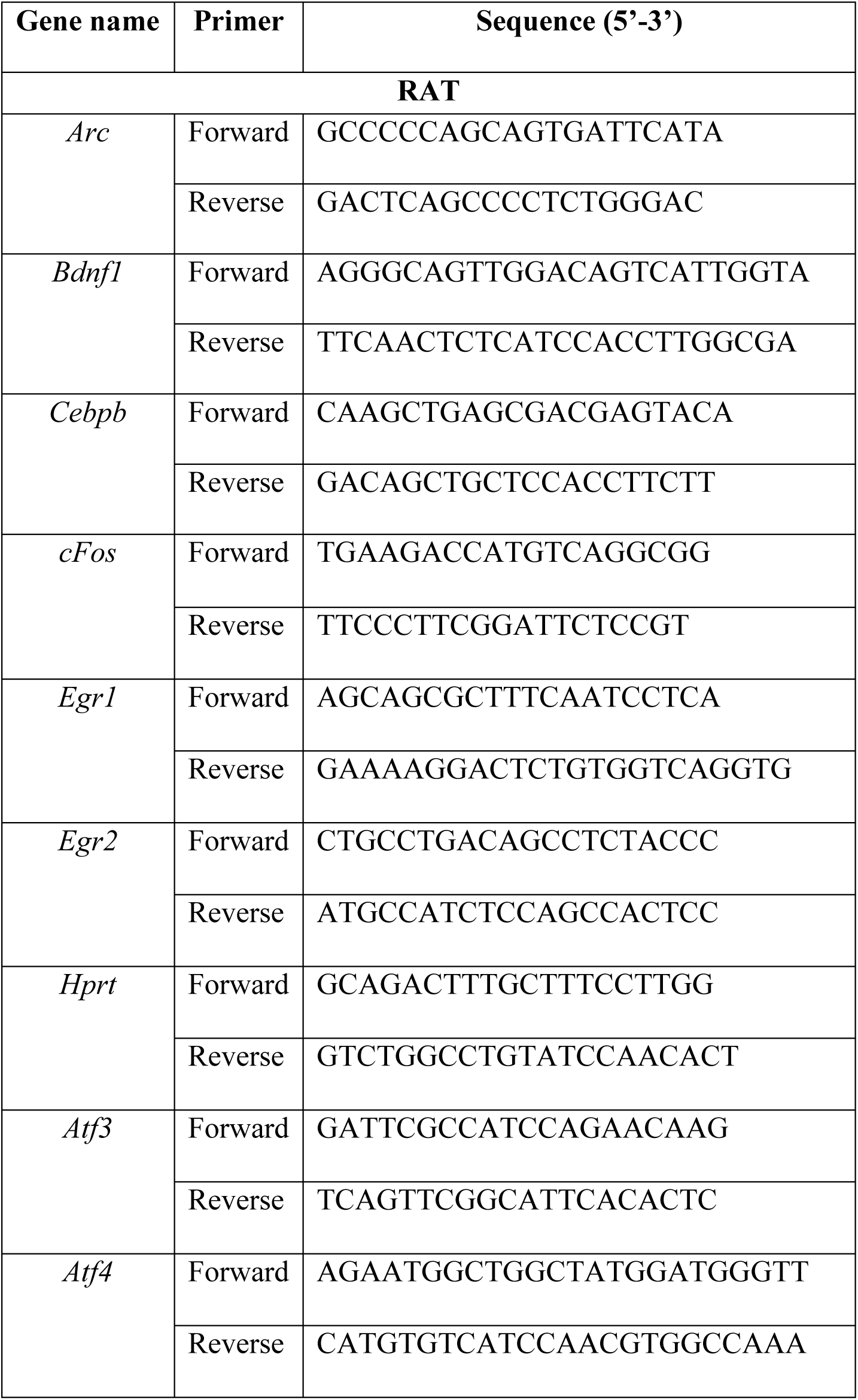
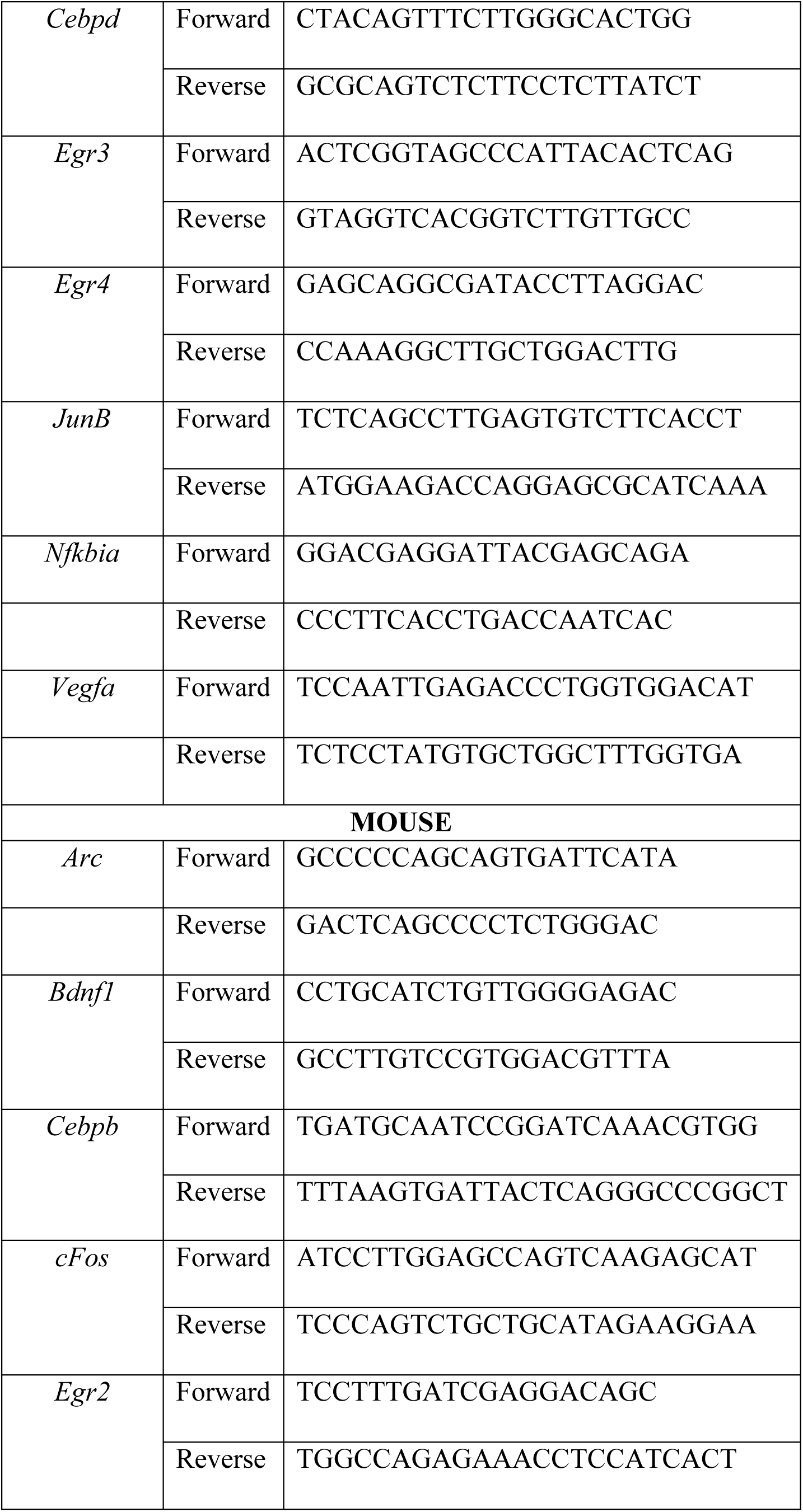
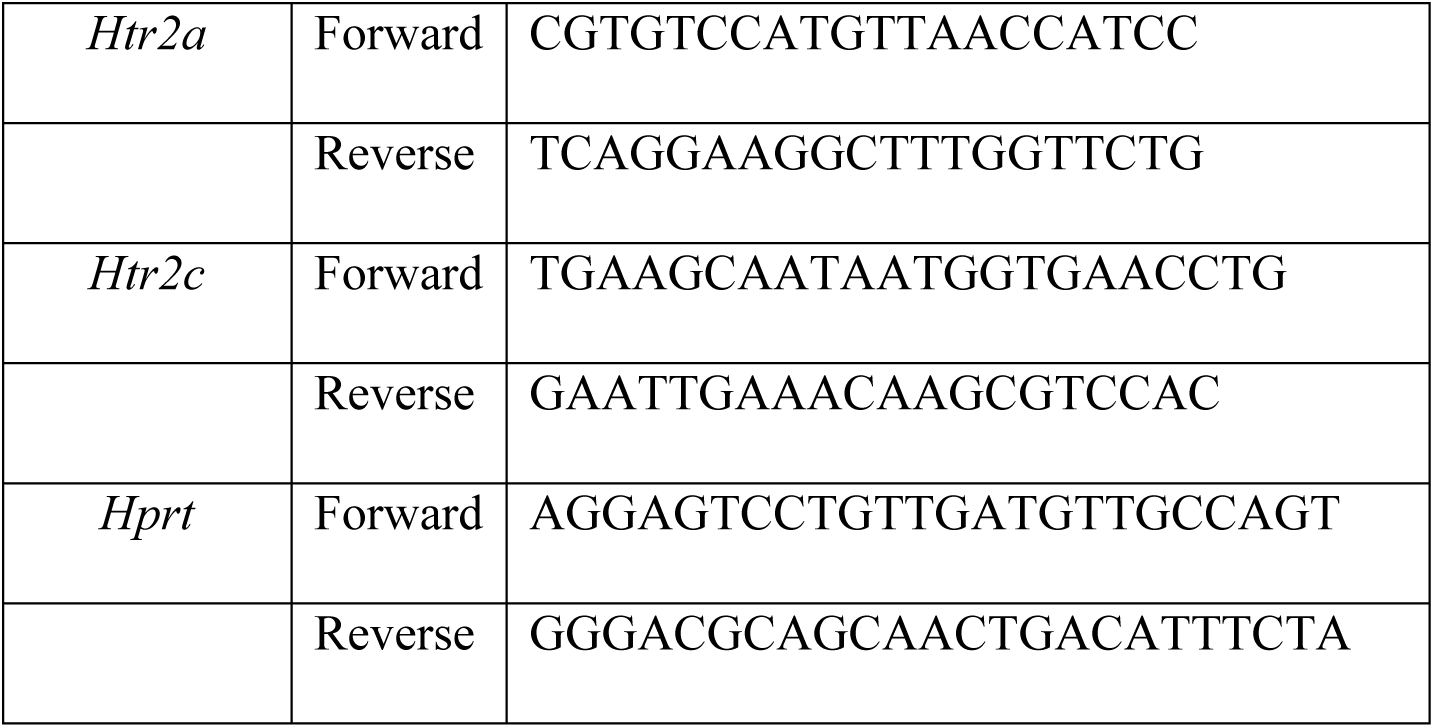
Primer sequences used for quantitative PCR (qPCR) analysis of rat/mouse cDNA

**Supplementary Table 2:**
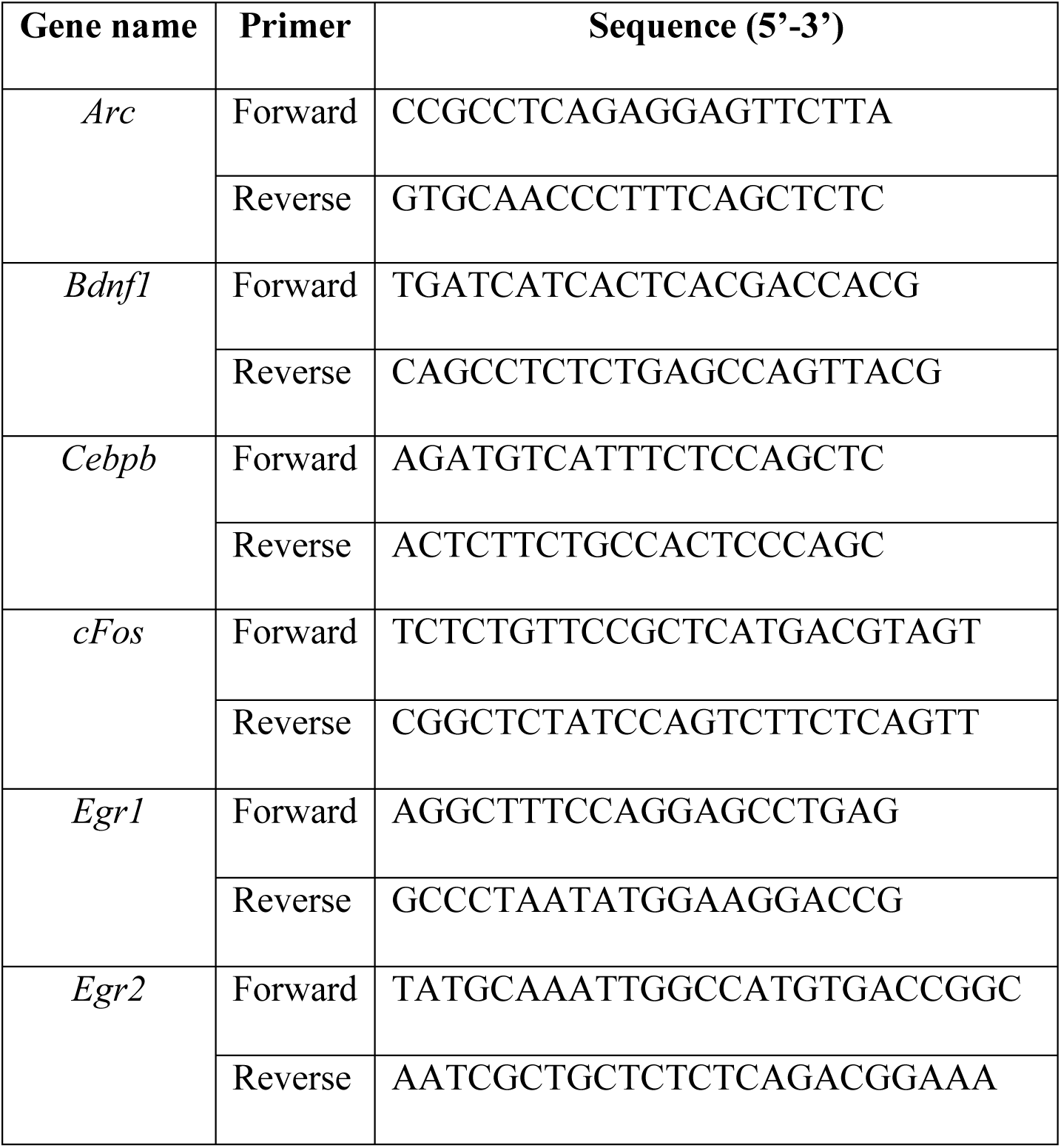
pCREB ChIP-qPCR primer sequences

